# Antigenic determinants of HAstV-VA1 neutralization and their relevance in the human immune response

**DOI:** 10.1101/2024.03.05.583477

**Authors:** Inci Ramírez-Bello, Tomás López, Rafaela Espinosa, Anisa Ghosh, Kassidy Green, Lidia Riaño-Umbarila, Carlos Gaspar-Castillo, Celia M. Alpuche-Aranda, Susana López, Rebecca M. DuBois, Carlos F. Arias

**Author notes:** Corresponding author: Carlos F. Arias.

## Abstract

Astroviruses are highly divergent and infect a wide variety of animal hosts. In 2009, a genetically divergent human astrovirus (HAstV) strain VA1 was first identified in an outbreak of acute gastroenteritis. This strain has also been associated with fatal central nervous system disease. In this work, we report the isolation of three high-affinity neutralizing monoclonal antibodies (Nt-MAbs) targeting the capsid spike domain of HAstV-VA1. These antibodies (7C8, 2A2, 3D8) were used to select individual HAstV-VA1 mutants resistant to their neutralizing activity and also select a HAstV-VA1 triple mutant that escapes neutralization from all three Nt-MAbs. Sequencing of the virus genome capsid region revealed escape mutations that map to the surface of the capsid spike domain, define three potentially independent neutralization epitopes, and help delineate four antigenic sites in rotaviruses. Notably, two of the escape mutations were found to be present in the spike sequence of the HAstV-VA1-PS strain isolated from an immunodeficient patient with encephalitis, suggesting that those mutations arose as a result of the immune pressure generated by the patient’s immunotherapy. In accordance with this observation, human serum samples exhibiting strong neutralization activity against wild-type HAstV-VA1 had a 2.6-fold reduction in neutralization titer when evaluated against the triple-escape HAstV-VA1 mutant, indicating shared neutralization epitopes between the mouse and human antibody response. The isolated Nt-MAbs reported in this work will help characterize the functional sites of the virus during cell entry and have the potential for developing a specific antibody therapy for the neurological disease associated with HAstV-VA1.

**Importance:** Human astroviruses (HAstVs) have been historically associated with acute gastroenteritis. However, the genetically divergent HAstV-VA1 strain has been associated with central nervous system disease. This work isolated high-affinity neutralizing monoclonal antibodies directed to HAstV-VA1. The proposed binding sites for these antibodies, together with previously reported sites for neutralizing antibodies against classical HAstVs, suggest the existence of at least four neutralization sites on the capsid spike of astroviruses. Our data show that natural infection with human astrovirus VA1 elicits a robust humoral immune response that targets the same antigenic sites recognized by the mouse monoclonal antibodies and strongly suggests the emergence of a variant HAstV-VA1 virus in an immunodeficient patient with prolonged astrovirus infection. The isolated Nt-MAb reported in this work will be helpful in defining the functional sites of the virus involved in cell entry and hold promise for developing a specific antibody therapy for the neurological disease associated with HAstV-VA1.

## Introduction

Human astroviruses (HAstVs), first identified in 1975, are significant etiological agents of infantile gastroenteritis and are also associated with diarrheal disease in the elderly and immunocompromised people (1, 2). The initially recognized HAstV strains were classified into eight serotypes currently known as classical HAstVs within the *Mamastrovirus 1* species in the *Mammastrovirus* genus of the *Astroviridae* family. Beginning in 2008, astrovirus strains, genetically closer to bovine, ovine, and porcine astroviruses rather than to classical HAstVs, were reported in human feces (3–5). These nonclassical HAstVs (MLB and VA) group into two different clades and are classified into three astrovirus species: *Mamastrovirus 6* (MLB1, MLB2, and MLB3), *Mamastrovirus 8* (VA2, VA4, VA5, and BF34), and *Mamastrovirus 9* (VA1 and VA3) (1). The association of viruses in the MLB and VA clades with gastroenteritis has not been conclusively demonstrated (1). However, they commonly infect humans, with a reported seropositivity rate of up to 100% in adult populations (6, 7).

More recently, HAstVs have been associated with meningitis and encephalitis in immunocompromised patients, with HAstV-VA1 (hereafter referred also as VA1) being the most frequently detected virus in these cases (see references in (8)). Also, a growing number of newly identified neurotropic astroviruses have been identified in cases of nonsuppurative encephalitis and neurological disease in pigs, ruminant species (cattle, sheep), and minks, changing the perception on the role of these viruses in non-enteric diseases (9). Mammalian astroviruses have high genetic diversity, are highly ubiquitous, and have a broad host range (10). Evolutionary analyses suggest sustained and extensive cross-species transmission events have occurred in the past between wild and domestic animal species and humans (10–13). In this context, studying human and animal astroviruses is warranted as part of a One Health approach and in preparation for future zoonotic events.

HAstVs are small, nonenveloped viruses with a single-stranded, positive-sense RNA genome of approximately 6.7 kb (1, 2). In the case of the nonclassical VA1 strain, the capsid of the mature, infectious virion is composed of two polypeptides: VP33, which constitutes the shell of the virus particle (basic region and inner core structural domain), and VP38, which forms dimeric globular spikes that protrude from the virion (outer core and spike structural domains) (14). These two polypeptides are derived from a capsid precursor protein of 86 kDa (VP86) processed intracellularly by an undefined protease (14). Previously, the antigenic sites of neutralizing monoclonal antibodies (Nt-MAbs) to the capsid spike domain of classical HAstVs have been mapped (15–17). However, despite the medical relevance of the non-classical HAstV-VA1, there is limited information about its antigenic sites. Its characterization should help to identify immunoreactive regions or epitopes as the basis of protective immunity and learn about functional sites on its capsid.

In this work, we report the isolation of three Nt-MAbs targeting HAstV-VA1. We identified mutations in the spike domain enabling the virus to escape neutralization by these MAbs, delineating three potentially independent neutralization sites. We also show that mouse and human neutralizing humoral immune responses target these antigenic regions. This study broadens our knowledge about the neutralization antigenic structure of HAstVs, including those more closely related to animal viruses, like VA1. It contributes to a more panoramic view of the functional sites of the virus capsid involved in the initial interactions of the virus with its host cell and provides the basis to develop a targeted antibody therapy for the neurological diseases associated with HAstV-VA1.

## Results

### HAstV-VA1 neutralization epitopes reside on the capsid spike domain

Previously, we reported that neutralization epitopes for classical human astroviruses serotypes 1, 2, and 8 are located on the capsid spike domain (16). To investigate if this is also the case for astrovirus HAstV-VA1, which belongs to a different clade than classical HAstVs, we evaluated the capacity of the core and spike structural domains of HAstV-VA1 to elicit virus-neutralizing antibodies (Nt-Abs). Rabbits were immunized with recombinant proteins corresponding to the core and spike structural domains, as well as a smaller region of the core domain (Fig. 1A). Antibodies directed to the virus capsid spike (aa 408 to 684 of ORF2; anti-Spike) efficiently neutralized the infectivity of HAstV-VA1 (Fig. 1B). In contrast, antibodies against two recombinant core proteins did not have neutralizing activity. These results strongly suggest that for HAstV-VA1, as in classical HAstVs, most neutralization epitopes reside on the capsid spike domain.

**Figure 1.**
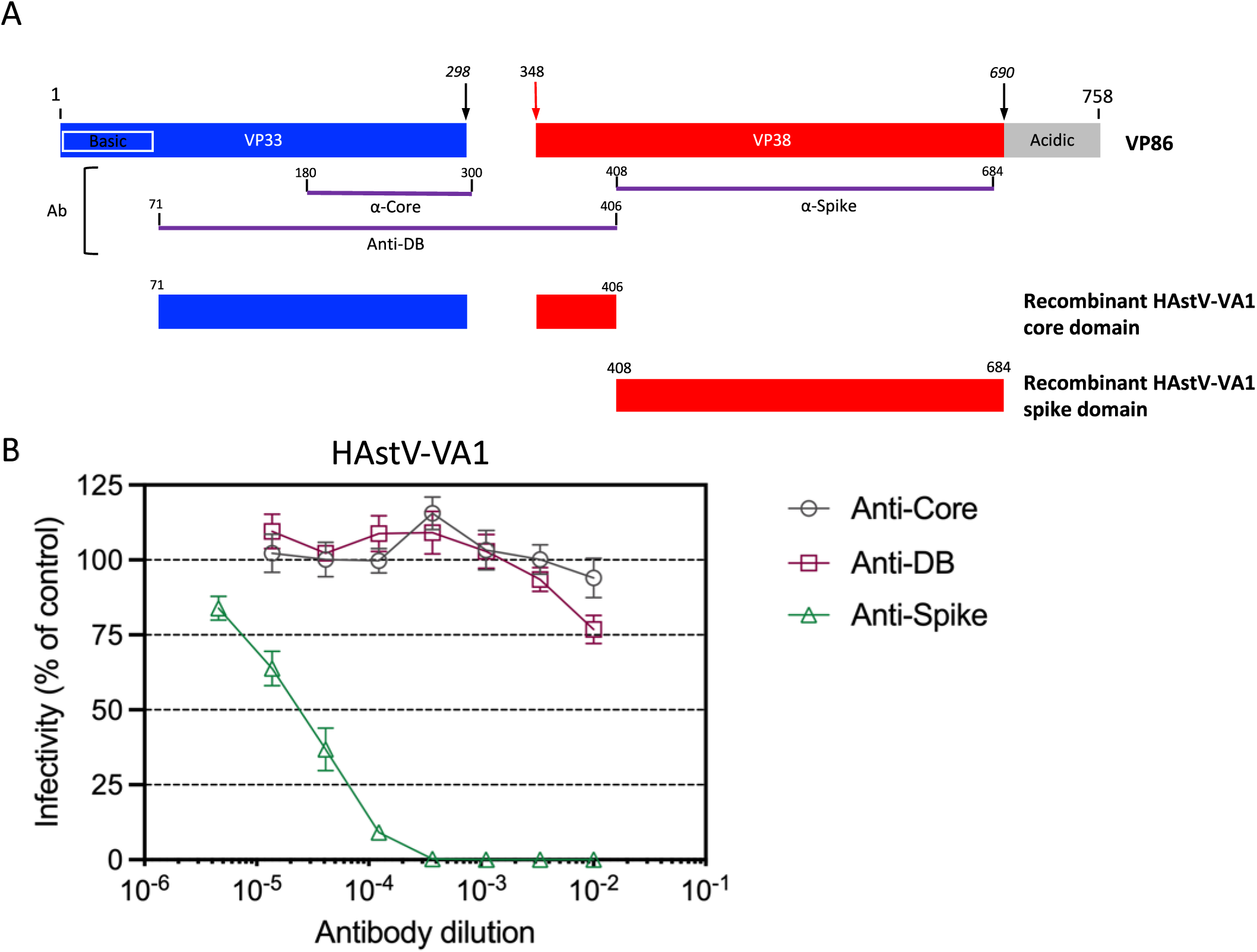
(A) Diagram of the VA1 ORF2 capsid precursor and recombinant proteins. ORF2 is represented as a box. The position of the core, spike, and acidic amino acid regions are indicated. The hatched box indicates a connector peptide. The recombinant proteins used to generate antibodies are represented as purple lines. The diagram is to scale. The numbers indicate the VA1 ORF2 amino acids included in each protein. (B) Induction of neutralizing antibodies by recombinant core, spike, and DB proteins. HAstV-VA1 was preincubated with rabbit anti-core or anti-DB or mouse anti-spike hyperimmune sera at the indicated dilutions, and the infectivity of the virus was determined as described in Materials and Methods. The infectivity assay was performed in biological triplicates and carried out in duplicate. The data are expressed as percentages of the value for the positive control (virus not incubated with antibodies) and represent the mean ± SEM.

### Isolation of neutralizing monoclonal antibodies to HAstV-VA1

To isolate Nt-MAbs against HAstV-VA1, BALB/C mice were immunized with HAstV-VA1 particles purified by CsCl isopycnic density centrifugation. Screening for hybridomas secreting antibodies to the virus was carried out by ELISA using purified virus particles as antigen (not shown), and positive hybridomas in this assay were subsequently tested in a neutralization assay. Three stable hybridomas secreting Nt-MAbs were identified: 7C8, 2A2, and 3D8. Ascitic fluids were produced in mice and evaluated for their neutralization activity to HAstV-VA1 (Fig. 2). All three MAbs efficiently neutralized virus infectivity at dilutions similar to those obtained with a mouse polyclonal antibody to the virus. In addition, they could efficiently detect infected cells by immunofluorescence (Fig. 2B) at 24 h post-infection, with most of the signal presumably corresponding to particles partially inside or at the edges of the vesicles induced during virus infection, as has been previously reported (18–20).

**Figure 2.**
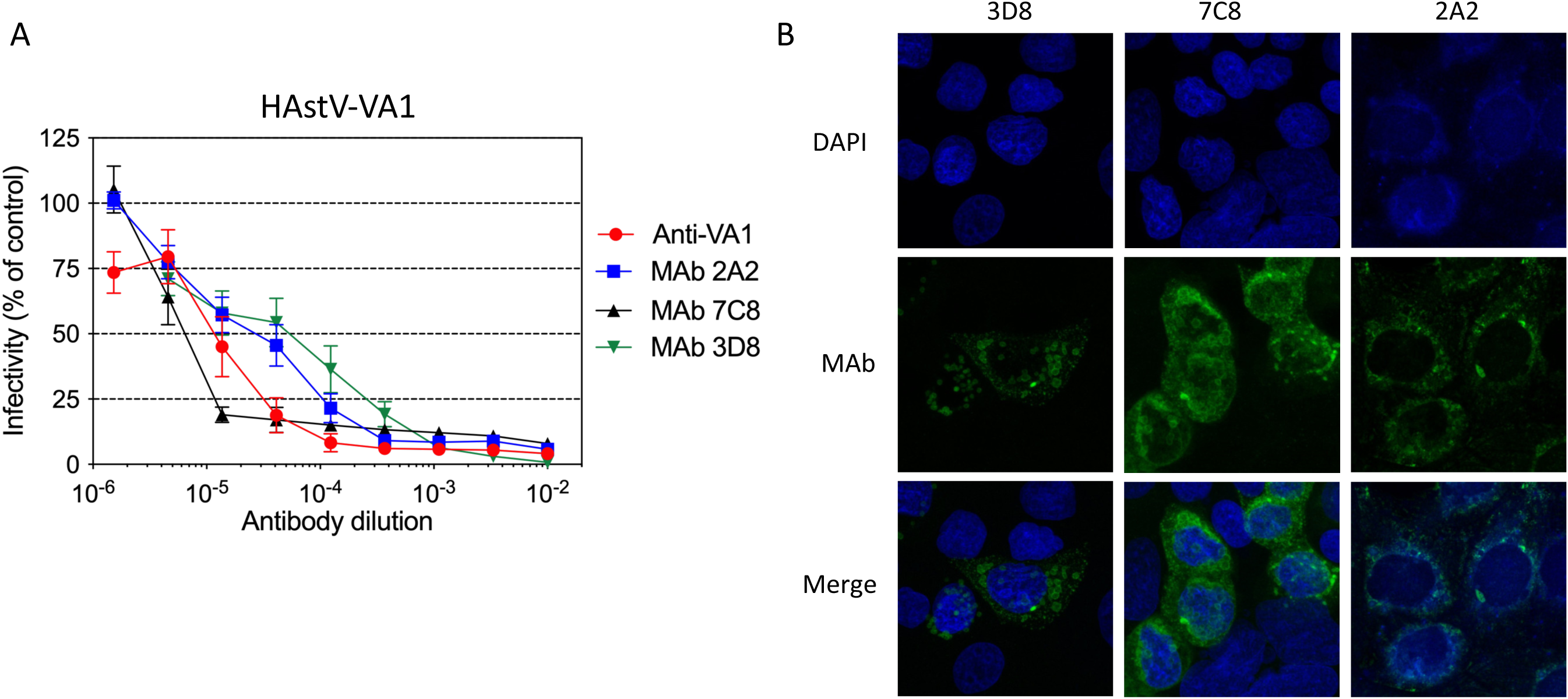
Neutralization activity of monoclonal antibodies. (A) HAstV-VA1 was preincubated with MAbs 2A2, 7C8, or 3D8 or mouse hyperimmune polyclonal serum to VA1 virus particles (Anti-VA1) at the indicated dilutions, and the infectivity of the virus was determined as described in Materials and Methods. The infectivity assay was performed in biological triplicates and carried out in duplicate, and the infectivity of the virus was determined as described in Materials and Methods. The data are expressed as percentages of the value for the positive control (virus not incubated with antibodies) and represent the mean ± SEM. (B) Caco-2 cells were infected with HAstV-VA1 at an MOI of 0.1, and at 24 hpi, they were fixed with formaldehyde and incubated with MAbs 2A2, 7C8, or 3D8. The nuclei were stained with DAPI. The cells were subsequently incubated with Alexa 488-labeled anti-IgG antibodies and observed for immunofluorescence as described in Material and Methods.

### Neutralizing antibodies to HAstV-VA1 have a high affinity for the capsid spike

The affinity of MAbs 7C8, 2A2, and 3D8 for recombinant HAstV-VA1 capsid spike was determined by biolayer interferometry. Anti-Mouse IgG Fc Capture (AMC) biosensors coated with MAb 7C8, 2A2, or 3D8 were dipped into wells containing 1:2 serially diluted HAstV-VA1 spike in assay buffer to determine the association rate. Biosensors were then dipped in assay buffer to determine the dissociation rate. We found that all three MAbs bind with high-affinity to the HAstV-VA1 spike, with dissociation constants (KD) of 1.2 nM for MAb 2A2, 0.4 nM for MAb 7C8, and 1.9 nM for MAb 3D8 (Fig. 3A to 3C).

**Figure 3.**
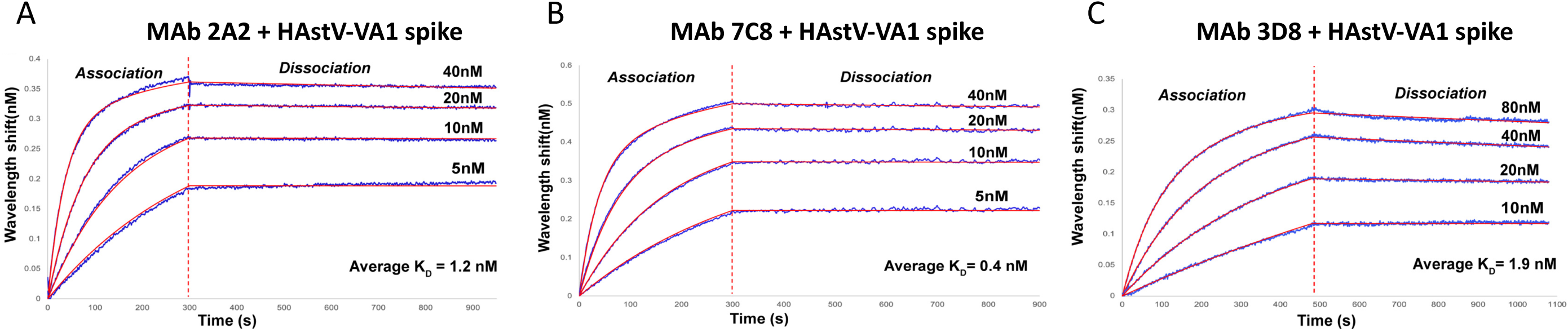
Binding affinities of mAbs 7C8, 2A2, and 3D8 to HAstV-VA1 capsid spike. (A) mAb 7C8, (B) 2A2, or (C) 3D8 was loaded onto biosensors and then placed into 1:2 serial dilutions of HAstV-VA1 spike at the indicated concentrations to allow association and then placed into assay buffer to allow dissociation. Representative traces of the data are shown. The KD value represents the mean of two independent experiments.

### Selection of HAstV-VA1 mutants that escape MAb neutralization

To determine the potential binding sites for the isolated Nt-MAbs, virus mutants that escaped neutralization were isolated after at least three virus passages in the presence of the corresponding Nt-MAb. As shown in Fig. 4, mutants that efficiently escaped neutralization by the cognate Mab were selected, although escape mutant m3D8 was still somewhat sensitive to neutralization at high MAb 3D8 concentrations. All three escape mutants were neutralized by the other two MAbs, suggesting that the three MAbs recognize different epitopes on the virus spike. In addition, all three escape mutants were neutralized similarly by the polyclonal antibody to HAstV-VA1.

**Figure 4.**
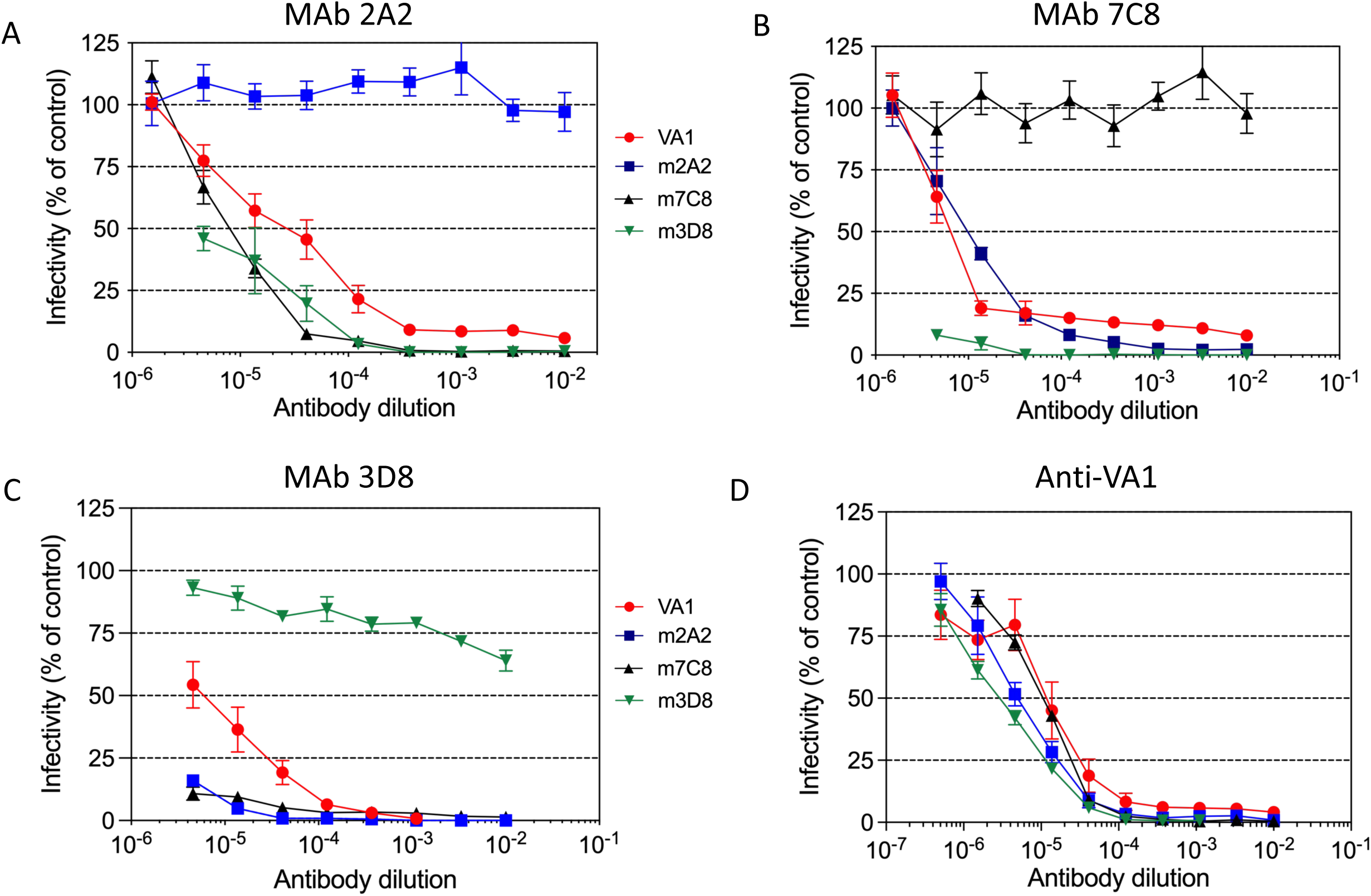
Neutralization of HAstV-VA1 escape variants. Escape variants m2A2, m7C8, and m3D8 or with wt HAstV-VA1 were preincubated with MAbs 2A2 (A), 7C8 (B), or 3D8 (C), or with mouse anti-VA1 polyclonal serum (D) at the indicated dilutions, and the infectivity of the virus was determined as described in Materials and Methods. The infectivity assay was performed in biological triplicates and carried out in duplicate. The data are expressed as percentages of the value for the positive control (virus not incubated with antibodies) and represent the mean ± SEM.

A double escape mutant, resistant to neutralization by both 7C8 and 2A2 MAbs, was isolated. This double mutant (m7C8/m2A2) was sensitive to neutralization by MAb 3D8 and by the polyclonal antibody to HAstV-VA1, confirming the independence of at least three neutralization antigenic sites on the virus spike (Fig. 5A). A virus mutant that escaped neutralization by all three Nt-MAbs, starting from the 7C8/2A2 double mutant and selecting a virus resistant to neutralization by MAb 3D8 was also isolated. Interestingly, this triple mutant was fully resistant to 3D8 neutralization and maintained its total resistance to Mab 7C8; however, it became slightly sensitive to neutralization by MAb 2A2 (Fig. 5B), indicating that the mutation selected by MAb 3D8 somehow facilitated the interaction of this triple mutant with MAb 2A2. It is important to point out, however, that 50% inhibition of the triple mutant by MAb 2A2 was reached at about a 1:300 dilution of the 2A2 ascites fluid (Fig. 5B). In comparison, 50% inhibition of the wild-type virus with the 2A2 ascitic fluid was obtained with about a 1:50,000 dilution (Fig. 2A), showing that the triple mutant was still very refractory to neutralization by all three Nt-MAbs. Finally, VA1 polyclonal serum was still effective in neutralization of the triple-escape mutant, suggesting that there is at least a fourth neutralization epitope on the HAstV-VA1 spike, different from the epitopes recognized by the three Nt-MAbs characterized in this work.

**Figure 5.**
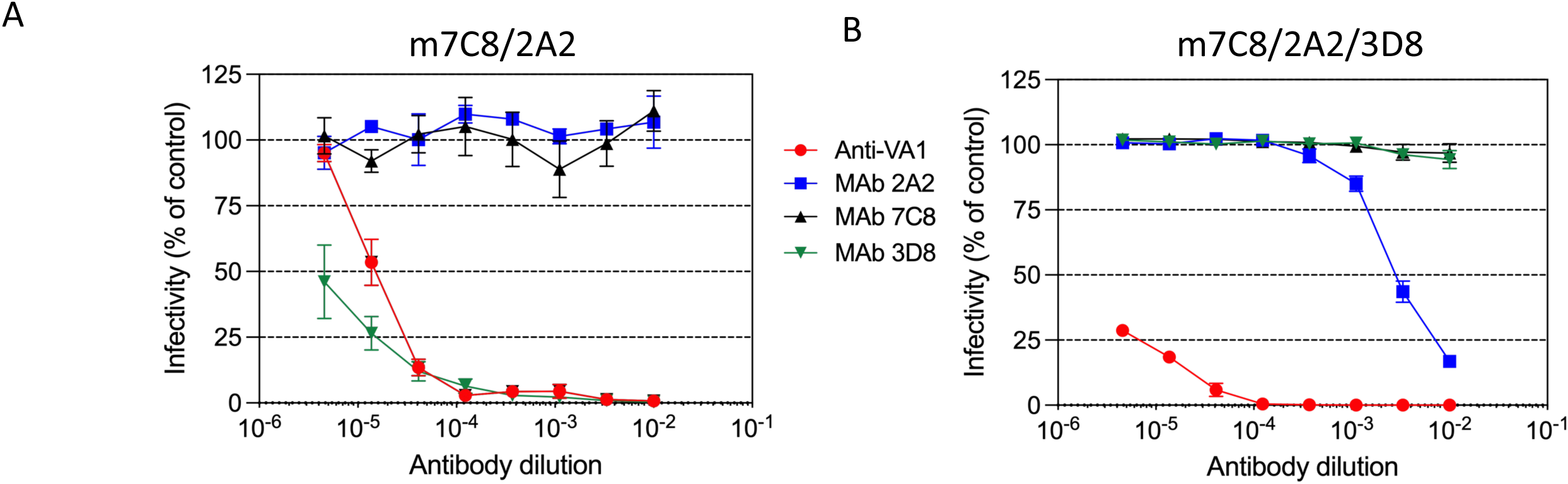
Neutralization of double and triple neutralization escape HAstV-VA1 variants. Double m7C8/2A2 (A) and triple m7C8/2A2/3D8 (B) neutralization escape variants were preincubated with MAbs 2A2, 7C8, or 3D8 or with mouse anti-VA1 polyclonal serum at the indicated dilutions, and the infectivity of the virus was determined as described in Materials and Methods. The infectivity assay was performed in biological triplicates and carried out in duplicate. The data are expressed as percentages of the value for the positive control (virus not incubated with antibodies) and represent the mean ± SEM.

### Mapping the mutations that allow HAstV-VA1 to escape mAb neutralization

To determine the amino acid changes that conferred HAstV-VA1 resistance to neutralization by the isolated MAbs, the sequence of the ORF2 region encoding the capsid spike of the wild-type and mutant viruses was determined. The spike sequence corresponding to the wild-type virus in our lab (HAstV-VA1, P14; GenBank #PP236967) has two conservative changes (Y420F and S670T) with respect to the sequence previously reported (GenBank #KY933670). When the P14 VA1 spike wild-type sequence was compared with those obtained from the escape mutants, a single amino acid change was detected on the VA1 mutants m7C8 (P543L) and m3D8 (E564V) spike (Fig. 6A). In contrast, the escape mutant m2A2 showed two amino acid changes separated by four amino acids (S611L and R615G) (Fig. 6A).

**Figure 6.**
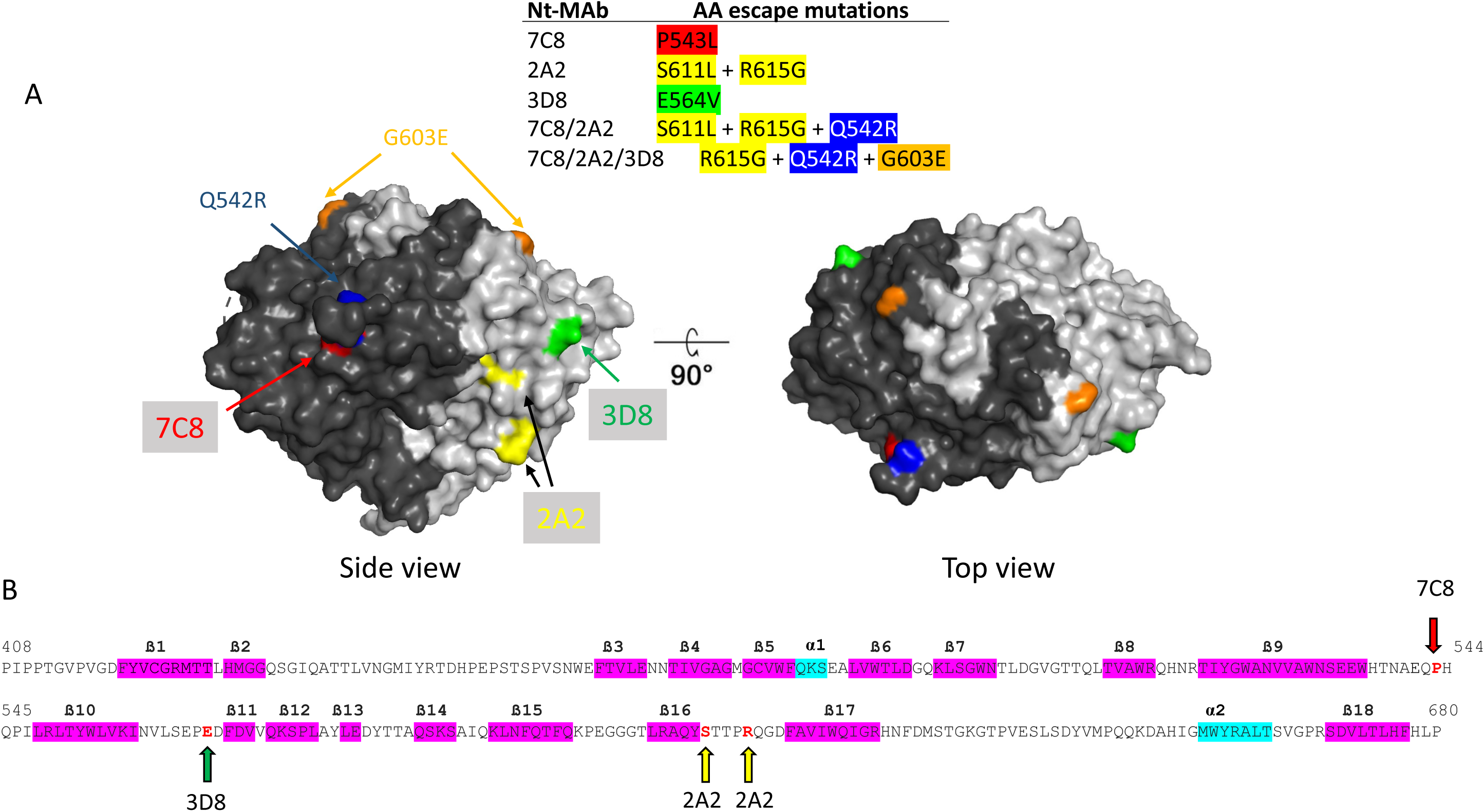
Localization of point mutations that confer escape from neutralization by monoclonal antibodies. (A) The crystal structure of the HAstV-VA1 spike is shown from the top and the side. One half of the dimer is dark gray, and the other half is light gray. The mutations that allow wt HAstV-VA1 to escape neutralization from the various MAbs are colored green for 3D8, yellow for 2A2, and red for 7C8. The additional mutations identified in the double m7C8/2A2 and triple m7C8/2A2/3D8 escape mutants are in blue and gold, respectively. The mutations are shown in only one protomer in the dimer. (B) Sequence of the HAstV-VA1 capsid spike protein (P14 strain). The amino acids that changed in the escape variants are in red and indicated by red (7C8), green (3D8), or yellow (2A2) arrows. Beta strands are highlighted in magenta, and alpha-helices in cyan.

In the 7C8/2A2 double escape mutant (m7C8/2A2), it showed the S611L and R615G mutations identified in m2A2, but instead of the P543L mutation detected in m7C8, the mutation Arg to Gly in the neighboring amino acid 542 (R542G) was selected. Finally, the triple escape mutant (m7C8/2A2/3D8) shared R615G and Q541R with the double mutant but acquired a new mutation at amino acid 603 (G603E) and lost mutation S611L. These results show the consistency of identification of sites that appear relevant for the Nt-MAbs interactions.

### Mapping of escape mutations on the 3D structure of the HAstV-VA1 spike

The 3D structure of the HAstV-VA1 spike has been recently reported (8). All escape mutations were found to be located on the spike surface (Fig. 6A), mapping to the β9-β10 loop (7C8), β10-β11 loop (3D8), and β16-β17 loop (2A2) (Fig. 6B). Escape mutations in viruses m2A2 and m3D8 are located close in space, on one of the faces of the spike. Thus, even though these two mutations appear to be part of two different neutralization epitopes, they could overlap and belong to the same antigenic site. On the other hand, the escape mutation in m7C8 maps in the middle region of the same face of the dimeric spike where 3D8 and 2A2 mutations lie, but in the opposite protomer (Fig. 6A). Of interest, the amino acid mutation selected by MAb 3D8 (G603E) in the triple escape mutant mapped close to the top of the spike, but not far from the initial escape mutation in m3D8 (E564V).

### The escape mutations in HAstV-VA1 can also be selected during virus evolution in a prolonged human VA1 infection

It was recently reported that the spike sequences of the characterized HAstV-VA1 strains can be classified into two distinct groups One group includes the spike sequences of viruses isolated from patients with gastrointestinal disease (VA1-gastro), while the second clade contains sequences from viruses isolated from immunocompromised people with neurological disease (VA1-neuro) (8). The recombinant spike domains for strain PS, a VA1-neuro strain (GenBank # ADH93577.1; ref 21), and strain P14, a VA1-gastro strain (GenBank # #PP236967) were generated in *E. coli*. Surprisingly, MAb 7C8 efficiently bound the P14 spike but not the PS spike by BLI (data not shown). Analysis of the spike sequence of the PS VA1-neuro strain showed 21 amino acid changes when compared to the P14 VA1-gastro strain (Fig. 7). Most interestingly, two contiguous changes in the PS VA1-neuro spike sequence are precisely the escape mutations identified in m7C8 and m7C8/2A2 escape viruses (Q542R and P543L) (Fig. 7), probably explaining the lack of interaction of MAb7C8 with the PS spike. These findings suggest that these amino acid changes arose in the PS VA1-neuro virus as a response to the imposed neutralizing immune pressure by the immunotherapy with human convalescent sera that the patient received (21).

**Figure 7.**
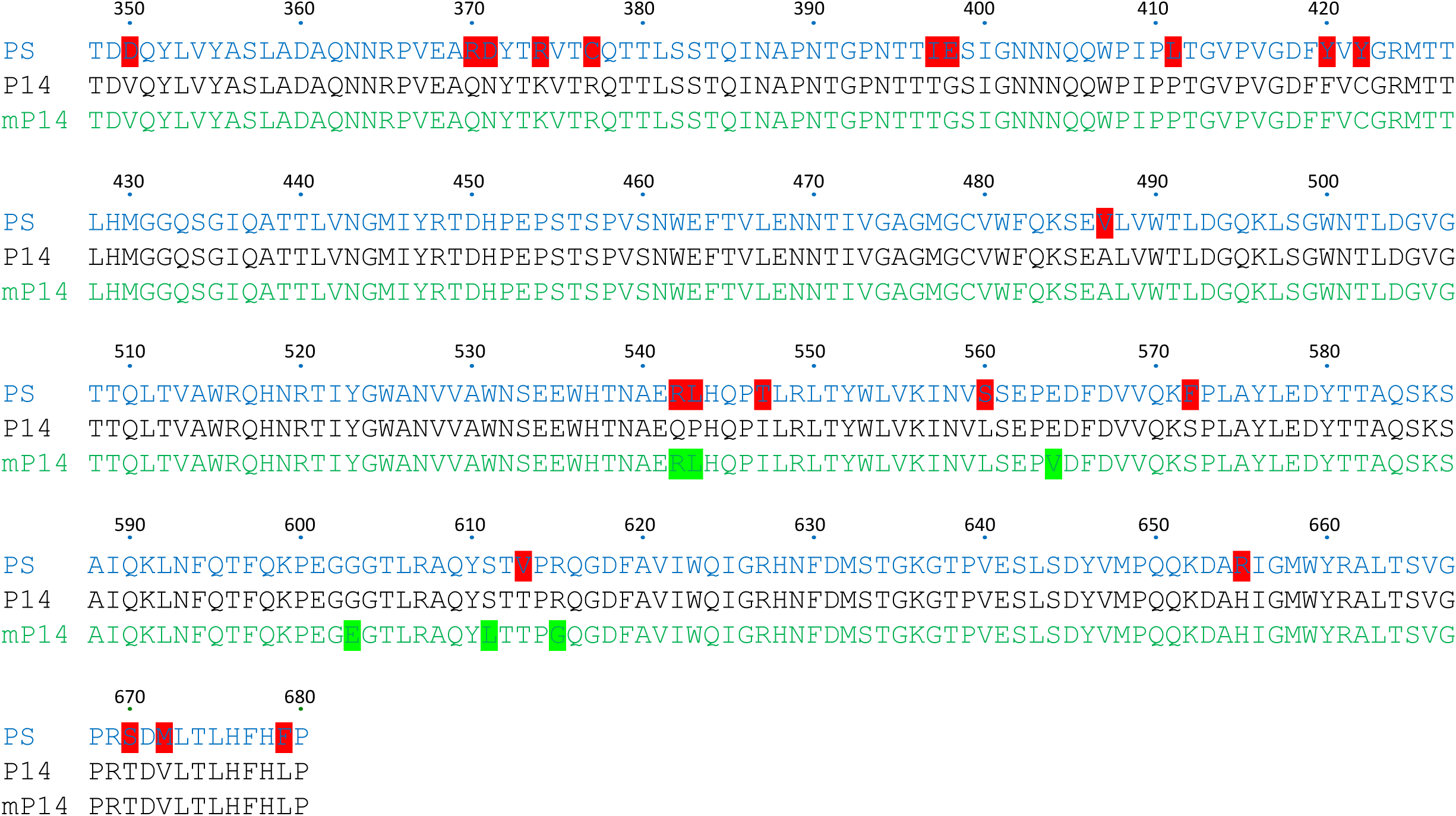
Comparison of the spike sequence of the PS (blue) and P14 (black) HAstV-VA1 strains. Amino acid differences are highlighted in red in the PS sequence. A composite of all escape mutations described in this work (mP14) is in green lettering. The mutations are highlighted in green.

### The neutralization sites identified in HAstV-VA1 are also targets of Nt-Abs induced during a natural HAstV-VA1 infection

Considering the previous observation, we evaluated if the neutralization sites identified in HAstV-VA1 with the Nt-MAbs described in this work are also targets for neutralizing antibodies induced during natural infection of HAstV-VA1 infection in humans. For this, an ELISA was used to test binding to the recombinant P14 VA1-gastro spike by human IgG antibodies present in 90 serum samples collected from healthy persons (mean age 40 years; range 6 to 60 years) who participated in different population-based serosurveys conducted in Mexico. Since no *bonafide* HAstV-VA1-negative human sera were available to be used as controls for the study, we provisionally considered those sera with OD values > 0.5 as positive for anti-VA1 antibodies (Fig. 8A). The ELISA data revealed that 72% (95%CI: 61.78, 81.15) of the samples showed OD values above 0.5, consistent with previous human serological studies showing high seroprevalence for HAstV-VA1 (7) (Fig. 8A). From this set of samples, we selected eight with the highest OD values and two with the lowest OD values (Fig. 8A). The antibody neutralization titer of these sera was then determined against the wild-type VA1-P14 strain and the triple-escape mutant VA1-P14 by a focus-reduction assay, showing that neutralization titers ranged from <1:200 to 1:51,200 (Fig. 8B). Of interest, the neutralization titers observed for wild-type VA1 with the different sera were reduced on average 2.6-fold when they were determined against the triple escape mutant VA1. These results suggest that the neutralization sites identified in the HAstV-VA1 spike with the mouse MAbs isolated in this study are also the target for the human neutralizing immune responses during natural infection with this virus.

**Figure 8.**
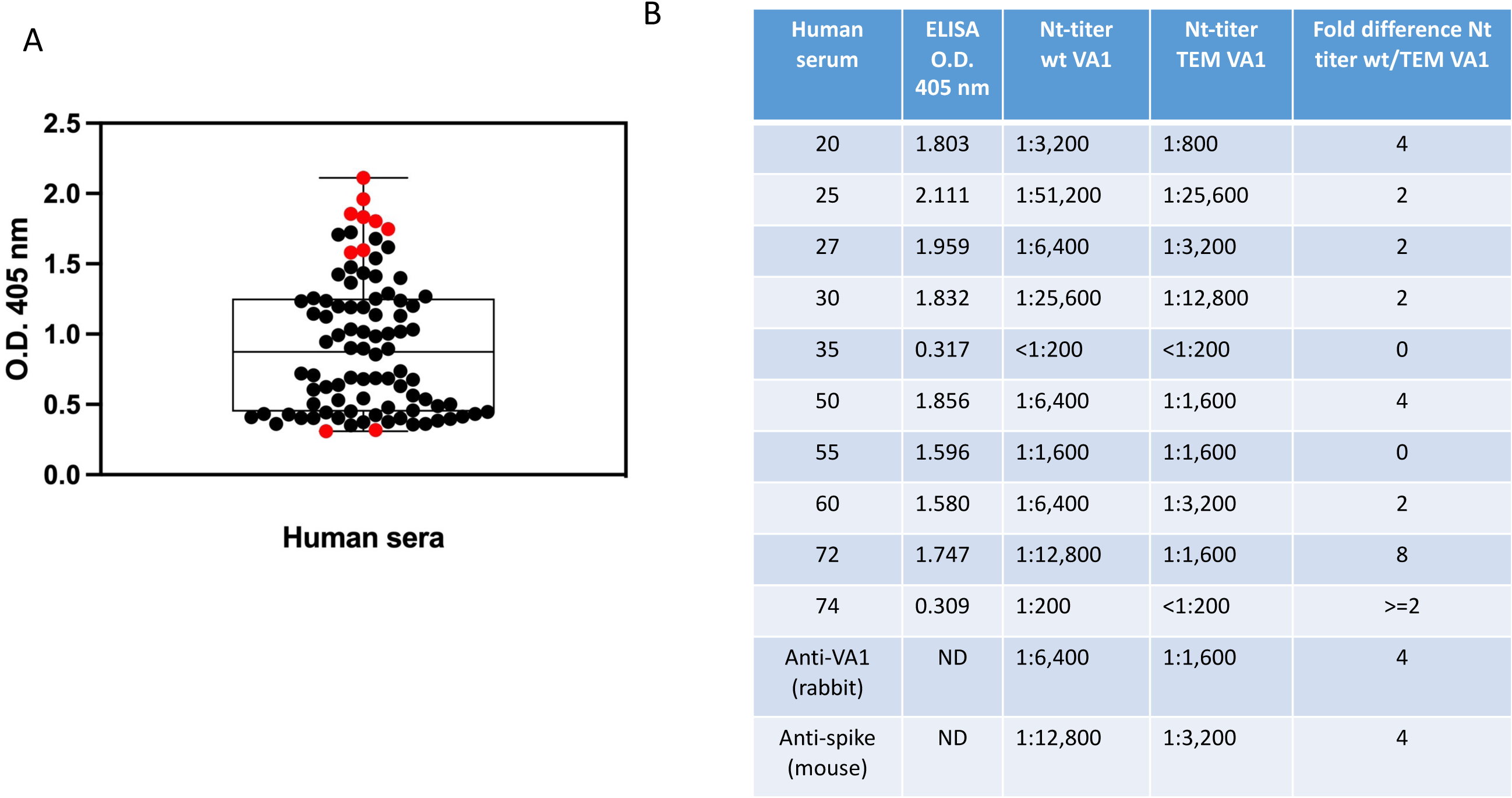
Titers of total and neutralizing human antibodies to HAstV-VA1. (A) The P14 recombinant spike was immobilized on ELISA plates and then incubated with a 1:100 dilution of a set of 90 human sera. The interaction was determined as described in Materials and Methods, and the optical density of the developed color was determined at 405 nm (O.D. 405). Each dot represents an individual serum, and the red dots represent the sera further analyzed by neutralization. The experiment was performed as two biological duplicates carried out in duplicate. The data are shown as a box and whisker plot. (B) Wild-type and triple escape mutant (TEM) HAstV-VA1 were incubated with a 1:100 dilution of human sera or rabbit and mouse polyclonal serum, and the infectivity of the virus was determined as described in Materials and Methods. The infectivity assay was performed in biological triplicates and carried out in duplicate. The neutralization titers are the serum dilution that neutralizes 50% of the infectious foci.

## Discussion

We have previously identified five neutralization epitopes on the virus spike of classical HAstV through mapping antibody escape mutations (16). All of these mutations were subsequently shown to lay in the epitope footprint of the cognate Nt-MAb by X-ray crystallography and cryoelectron microscopy analyses of the antibody-spike complexes (17; unpublished data), suggesting that escape mutations can indeed be a proxy reference site for the Nt-MAb binding epitopes on the astrovirus spike domain. In this work, we have isolated and mapped mutations that allow HAstV-VA1 to escape neutralization by Nt-MAbs. As in classical HAstVs, the escape mutations are located on the capsid spike domain of HAstV-VA1. In agreement with this observation, polyclonal antibodies elicited by the recombinant spike domain, but not those elicited by the core domain, elicited high titers of Nt-Abs (Fig. 1B), as previously reported for classical HAstV (16).

The HAstV-VA1 Nt-MAb escape mutations identified three independent neutralization epitopes that appear to be located in two different antigenic sites, or regions that contain overlapping epitopes recognized by different antibodies. One antigenic site would be represented by the binding site of MAb 7C8, and the second by MAbs 2A2 and 3D8. These two sites are present on the same face of the spike but on opposite spike protomers. However, it is important to remember that some escape mutations could affect MAb neutralization indirectly through larger structural alterations to prevent its binding to the virus.

Classical and VA viruses represent two different HAstV clades. However, although their spike proteins have low sequence identity and apparent structural differences (8), they share a folding topology and a dimeric structure, which is also maintained in the MLB1 spike (22) and the recently described spike structure of murine astrovirus (23). In this regard, the previously reported escape mutations to Nt-Abs directed to classical HAstV serotypes 1, 2, and 8 (16) seem to lay in regions similar to those described in this work for the HAstV-VA1 spike (Fig. 9A and 9B). Thus, the neutralization epitope potentially recognized by the HAstV-VA1 Nt-MAb 7C8 seems to lay, topologically, very close to the epitope recognized by Nt-MAb 2D9 on the HAstV-8 spike, probably defining a similar antigenic site. Also, the epitopes recognized by MAbs PL2 and 3E8 on HAstV-2 and HAstV-8, respectively, represent, topologically, a similar antigenic site to that defined by HAstV-VA1 MAbs 2A2 and 3D8. In addition, MAbs 3B4 and 4B6, to HAstV-1 and HAstV-2 appear to form an additional antigenic site located at the top of the spike. Finally, and very interestingly, the epitope recognized by MAb 3H4 to HAstV-1 appears to represent an entirely different neutralizing antigenic site, located in the lower region of the spike (Fig. 9B). Altogether, the neutralization escape mutations in both classical and HAstV-VA1 virus suggest the existence of at least four neutralization antigenic sites on the astrovirus spike, located in either of the two protomers and the lower, middle or top of the dimeric structure, highlighting the vulnerability of astrovirus to neutralization by the antibody immune response.

**Figure 9.**
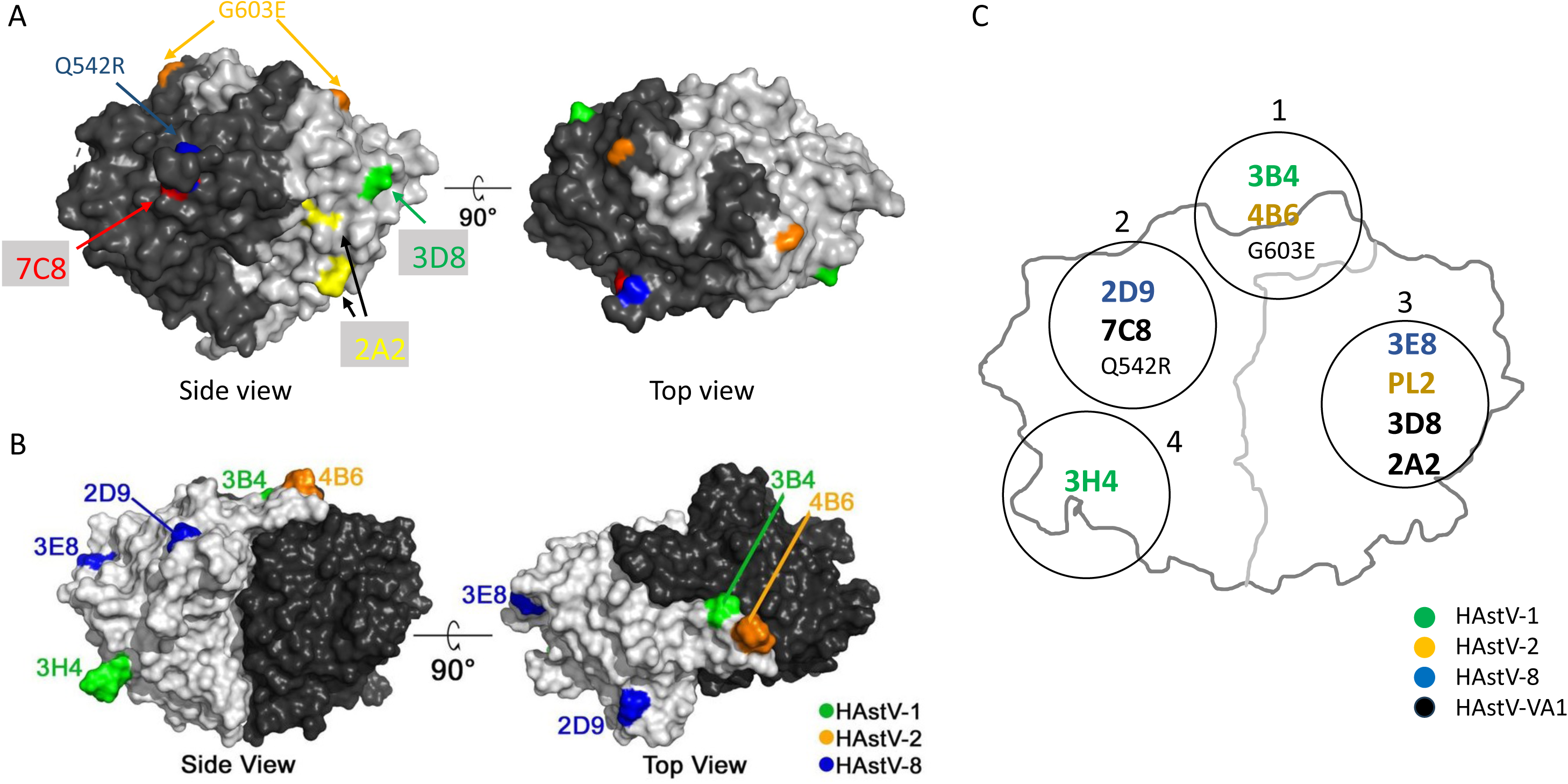
(A) Crystal structure of the HAstV-VA1 spike domain. The mutations that allow wt HAstV-VA1 to escape neutralization from the various MAbs are shown in different colors. The additional mutations identified in the double m7C8/2A2 and triple m7C8/2A2/3D8 escape mutants are in blue and gold, respectively. (B) The mutations that allow HAstV-1, −2, and −8 to escape neutralization (see (16)) are indicated in the crystal structure of the HAstV-2 spike domain. Both crystal structures are shown from the top and the side. One-half of the dimers is dark gray, and the other half is light gray. In both structures, the mutations are shown in only one protomer in the dimer. (C) Diagrammatic scheme of the HAstV-2 spike domain. The circles represent the proposed antigenic domains, with the various Nt-MAb escape mutations indicated inside the circles.

Of interest, two sites conserved on the spike of the eight classical HAstVs have been proposed as potential binding sites for a cell receptor (24). One of them, the P site located on the side of the spike, is in a similar location to that of the conserved C site identified when 15 different mouse astrovirus spike sequences were compared (23). These topologically conserved sites in classical human and murine astroviruses seem to overlap with the antigenic site 2 proposed in this work (Fig. 9C). On the other hand, the conserved S site in classical HAstVs (24) appears to overlap with antigenic site 4 (Fig. 9C). These regions may represent functional sites conserved among different astrovirus species.

In 2010, the spike sequence of a HAstV-VA1 strain (Genbank accession# ADH93577.1) was reported, originating from a frontal cortex biopsy specimen (21). This specimen was obtained from a 15-year-old boy with agammaglobulinemia who developed fatal astrovirus encephalitis was reported. Notably, the patient received monthly intravenous immunoglobulin therapy during his hospitalization. The amino acid sequence of the spike domain of this virus, HAstV-VA1-PS, has 21 amino acid changes compared to the sequence of the HAstV-VA1-P14 virus in our lab (Fig. 7). Remarkably, two of these changes, Q542R and P543L, which are otherwise conserved in all HAstV-VA1 capsid sequences reported thus far (8) are identical to escape mutations selected by Nt-MAb in this work. In addition, a second amino acid change in the VA1-PS spike, T613V, lies between amino acids 611 and 615, the two changes selected in the escape mutant to Nt-MAb 2A2 (Fig. 7). Altogether, these observations strongly suggest that the highly mutated VA1-PS strain, compared to the VA1-P14 strain, and in particular some of the mutations in the spike domain sequence, might have arisen due to the immune pressure imposed by the immunoglobulin therapy given to the patient. This driver has also been suggested for the emergence of the first SARS-CoV-2 variant of concern, Alpha (25, 26), and for the appearance of the original Omicron variant and its descendent BA.2.86, hyper-mutated variants, which most likely appeared in patients that received immunotherapy with convalescent plasma (27–29).

We also show here that the neutralization antigenic sites reported in this work can be the target of neutralizing antibodies in humans during natural infection with the VA1 virus, supporting a relevant functional role of these sites in the HAstV spike during infection of the susceptible epithelial gastrointestinal cells. Moreover, the finding that the neutralization antibody titer to HAstV-VA1 in a set of human sera (Fig. 8) had a 2.6-fold reduction in their neutralization titer when tested against the triple escape mutant of the VA1 virus suggests that the antigenic sites represented by MAbs 7C8, 2A2, and 3D8 are immunodominant during a natural human VA1 infection. It remains to be determined if these sites on the virus overlap with binding sites for the cell receptor or co-receptors. In this regard, it has recently been reported that the neonatal Fc receptor (FcRn) facilitates classical HAstV cell infection at a post-binding step (30). However, FcRn does not seem involved in HAstV-VA1 infection (30; unpublished data). The determination of the structure of the FcRn-HAstV spike complex and the effect of Nt-MAbs on the spike-FcRn interaction will help to understand the mechanism of neutralization of the reported Nt-MAbs to classical HAstVs. It should also give insights into the functional regions of nonclassical VA and MLB viruses.

The characterization of the neutralization mechanisms of the reported anti-VA1 Nt-MAbs, together with the identification of the actual epitopes of the antibodies on the virus spike by structural biology approaches, as has been previously shown for classical HAstV (17), will aid in identifying virus functional sites required for astrovirus cell entry. This information will be relevant for the development of prophylactic and therapeutic approaches to address this neurotropic astrovirus strain.

## Materials and Methods

### Ethical considerations

The samples were collected in three population-based serosurveys conducted in Tapachula, Chiapas (Sep 2018), Puente de Ixtla, Morelos (Jun 2016), and Campeche y Hopelchén, Campeche (Jun 2017). The participants were selected randomly, were older than two years, and had no fever or other clinical manifestation at the time of sampling. Patient blood sampling protocol was approved by the Research Committee of the Instituto Nacional de Salud Pública (protocol/approval number: 1498/CI-776-2017 and 1449/CI-279-2017). The authors state that all procedures contributing to this work comply with the ethical standards of the Helsinki Declaration of 1975, as revised in 2008.

### Cells and viruses

Caco-2 cells, clone C2Bbe1 (ATCC), were propagated in DMEM Advanced (Gibco No. 12491015) supplemented with 5% fetal bovine serum (FBS) (Cansera) and glutamine 2 mM in a 10% CO2 atmosphere at 37°C. HAstV-VA1 was obtained from David Wang (Washington University, St. Louis, Missouri). VA1 virus was purified by isopycnic gradient centrifugation as previously described (14).

### Recombinant proteins

Expression and purification of PS and P14 HAstV-VA1 spikes were carried out as previously described (8). Proteins containing amino acids 71 to 406 (anti-DB), 180-300 (anti-core), and 408 to 684 (anti-spike) of VA1 ORF2 (GenBank # NC_013060.1) were synthesized in E. coli and purified as previously reported (14).

### Antibodies

Recombinant proteins representing different regions of VA1 capsid precursor protein (described above; the spike protein sequence corresponds to the P14 strain) were used to generate hyperimmune sera in New Zealand rabbits, as reported previously (16). Rabbit and BALB/c mouse hyperimmune sera to VA1 were generated by immunization with either 250 ug (rabbits) or 50 ug (mice) of purified virus particles in Freund’s complete adjuvant followed by three immunizations every two weeks, with the same amount of virus in Freund’s incomplete adjuvant.

### Monoclonal antibodies isolation

Eight-week-old BALB/c mice were immunized with 50 ug of purified VA1 virus at 1:1 with Freund’s complete adjuvant, and three more immunizations were performed every two weeks using the same amount of virus in Freund’s incomplete adjuvant. Four days after the last immunization, the spleen was extracted, and splenocytes were fused with Fox myeloma cells using 50% polyethylene glycol; the cells were suspended in hypoxanthine-aminopterin-thymidine medium and directly plated in 96-well plates. Hybridomas secreting antibodies to HAstV-VA1 were screened by an ELISA (see below), and those positive by the ELISA were then assayed in a neutralization assay (see below). The hybridomas of interest were cloned three times by limiting dilution using peritoneal macrophage feeder layers. Selected MAbs were amplified as mouse ascitic fluid.

### ELISA

Hybridomas positive for monoclonal antibodies to VA1 were selected using purified VA1 virus particles, as described before, at a 2 ug/ml concentration in 96-well ELISA microtiter plates. Briefly, after virus adsorption, the plates were washed with PBS containing 0.1% Tween 20 and blocked with 1% BSA in PBS/Tween for 1 h at 37°C. Conditioned medium from hybridoma cells or mouse hyperimmune serum was added as a control and incubated at 37°C for 1 h. After washing the plates, they were incubated with goat anti-mouse antibody conjugated to alkaline phosphatase (KPL). The reaction was developed by adding the phosphatase substrate (Sigma 104; 1 mg/ml), and the absorbance was recorded at 405 nm. In the case of human sera, the above procedure was followed, except that the VA1 P14 spike was used as antigen, and the sera was tested at a 1:100 dilution. A secondary anti-human IgG coupled to peroxidase (Aviva SB) was added at a dilution of 1:1,000 in PBS.

### Immunofluorescence

Caco-2 cells grown on coverslips were infected with HAstV-VA1 at an MOI of 5. At 24 h post-infection, the cells were fixed with 2% paraformaldehyde in PBS for 20 min and permeabilized by incubation with 0.5% Triton X-100 in blocking buffer (1% bovine serum albumin in PBS with 50 mM NH4Cl) for 15 min. After washing with PBS, the coverslips were incubated with 2A2 (1:2,500), 7C8 (1:2,000), or 3D8 (1:1,000) ascitic fluids for 1 h at room temperature. The cells were then washed with PBS, and Alexa 488-labeled anti-IgG antibody (Invitrogen) was added at a 1:1,000 dilution for 1 h at room temperature. Nuclei were stained with 30 nM DAPI (4=,6-diamidino-2-phenylindole) for 15 min. Coverslips were mounted on glass slides by use of Citifluor AF100 antifade solution (Citifluor Ltd., London, United Kingdom), and the samples were observed under a Zeiss Axioskop 2 fluorescence microscope coupled to a digital camera (Photometrics Cool Snap HQ).

### Neutralization assays

The indicated concentration of ascitic fluid, hyperimmune sera, or human sera was preincubated at a multiplicity of infection (MOI) of 0.004 of wild-type, double- or triple-escape mutants of HAstV-VA1 strain for 1 h at room temperature. The virus-antibody mixture was then added to confluent C2Bbe1 cell monolayers grown in 96-well plates and incubated for 1 h at 37°C. After this time, the cells were washed three times with minimum essential medium (MEM) without serum, and the infection was left to proceed in DMEM-HG supplemented with non-essential amino acids for 24 h at 37°C. Infected cells were detected by an immunoperoxidase focus-forming assay, as described previously (14), but using the anti-VA1 polyclonal antibody diluted 1:2,000 for detecting the viral antigens. The neutralization antibody titer was defined as the antibody dilution that blocks at least 50% of the input virus.

### Biolayer interferometry experiments

Biolayer interferometry (BLI) data was collected on an Octet RED384 using the Data Acquisition Software (version 11.1.1.19), with the temperature set to 25 °C and shaking at 1000 rpm. BLI experiments were performed in assay buffer (PBS, 1% BSA, 0.05% Tween 20). Anti-Mouse IgG Fc Capture (AMC) biosensors were hydrated in assay buffer for at least 10 min before starting the experiment. BLI experiments were performed as follows:

(1) pre-hydrated AMC biosensors were dipped in assay buffer for 60 sec to establish a baseline;
(2) biosensors were dipped into a 1:200 dilution of mAb 7C8, 2A2, or 3D8 ascites fluid in assay buffer for 120 sec to load the mAb onto the biosensor; (3) biosensors were dipped into assay buffer for 60 s to confirm stable mAb loading and establish a new baseline; (4) mAb-loaded biosensors were dipped in wells containing 1:2 serially diluted HAstV-VA1 spike in assay buffer at the indicated concentrations for at least 300 sec to determine the association rate; (5) biosensors are dipped in assay buffer for 600 sec to determine the dissociation rate. Each curve was reference-subtracted with a 0 nM HAstV-VA1 spike control and aligned to the baseline and dissociation steps for inter-step correction. A global association 1:2 bivalent analyte model using the Octet Data Analysis HT software v7 (Sartorius) was used to account for the dimeric HAstV-VA1 spike. At least three curves were used to determine the on- and off-rate and calculate the dissociation constant (KD). Average KD values are reported as the mean of two independent experiments.

### Sequencing

The nucleotide sequences of the wild-type or mutant VA1 spike were determined as reported elsewhere (16) but using the following primers: 5-GTCCAATATCTAGTTTATG-3 (SpikeVA1up), corresponding to nucleotides 5259 to 5277 of HAstV-VA1 (accession number NC_013060), and 5-GGCGCAATTTTTTCTTGAC-3 (SpikeVA1lw), corresponding to nucleotides 6453 to 6471 of the virus. After PCR amplification, the product was sequenced using Sanger chemistry at the Instituto de Biotecnología, Universidad Nacional Autónoma de México sequencing facility.

### Isolation of neutralization escape variants

To select escape variants, we incubated viral lysates (with at least 10^7^ focus-forming units [FFU]/ml) with the appropriate MAb at a dilution of 1:100 to 1:1,000 of the ascitic fluid for 1 h at room temperature. The mix was then used to infect CaCo-2 monolayers grown in 6-well plates for 1 h at 37°C, and the unbound virus was removed by washing three times. The cell monolayers were incubated at 37°C for 72 to 96 h post-infection (hpi) in DMEM-HG supplemented with non-essential amino acids and tetracycline (1 μg/ml), and the corresponding MAb diluted 1:1,000. Viral lysates were then prepared by three freeze-thawing cycles, and the procedure was repeated at least three times before confirming the phenotype. The phenotype of the variant viruses was evaluated by a neutralization assay, and the selection was repeated until detection of the neutralization escape variant (this usually took 4 or 5 passages in the presence of the Nt-MAb). The double-escape variants were selected as described above, starting with the single variant instead of the wild-type virus.

### Statistics

Statistical analysis was determined by a two-tailed T-test with a 99% confidence interval using GraphPad Prism 9.0.1 Software (GraphPad Software, Inc.).

## Acknowledgments

This research was partially supported by grants NIH R01 AI144090 to R.M.D. and C.F.A., CONACyT 302965 to S.L.; and DGAPA IN210120 to T.L. We are grateful to David Wang (Washington University) for kindly providing human astrovirus VA1. We thank Marco Espinoza for his help in cell culture, Elizabeth Mata and Graciela Cabeza, and the animal house personnel for help during animal handling and immunization. We also thank Santiago Becerra Ramirez, Eugenio Lopez Bustos and Jorge Arturo Yañez Ponce de León for their help in the sequencing core facilities of the Instituto de Biotecnología.

